# Topconfects: a package for confident effect sizes in differential expression analysis provides improved usability ranking genes of interest

**DOI:** 10.1101/343145

**Authors:** Paul F. Harrison, Andrew D. Pattison, David R. Powell, Traude H. Beilharz

## Abstract

**Background:** A differential gene expression analysis may produce a set of significantly differentially expressed genes that is too large to easily investigate, so that a means of ranking genes by their biological interest level is desirable. The life-sciences have grappled with the abuse of *p*-values to rank genes for this purpose. As an alternative, a lower confidence bound on the magnitude of Log Fold Change (LFC) could be used to rank genes, but it has been unclear how to reconcile this with the need to perform False Discovery Rate (FDR) correction. The TREAT test of McCarthy and Smyth is a step in this direction, finding genes significantly exceeding a specified LFC threshold. Here we describe the use of test inversion on TREAT to present genes ranked by a confidence bound on the LFC, while still controlling FDR.

**Results:** Testing the Topconfects R package with simulated gene expression data shows the method outperforming current statistical approaches across a wide range of experiment sizes in the identification of genes with largest LFCs. Applying the method to a TCGA breast cancer data-set shows the method ranks some genes with large LFC higher than would traditional ranking by *p*-value. Importantly these two ranking methods lead to a different biological emphasis, in terms both of specific highly ranked genes and gene-set enrichment.

**Conclusions:** The choice of ranking method in differential expression analysis can affect the biological interpretation. The common default of ranking by *p*-value is implicitly by an effect size in which each gene is standardized to its own variability, rather than comparing genes on a common scale, which may not be appropriate. The Topconfects approach of presenting genes ranked by confident LFC effect size is a variation on the TREAT method with improved usability, removing the need to fine-tune a threshold parameter and removing the temptation to abuse *p*-values as a de-facto effect size.

## Background

Misunderstanding and abuse of *p*-values has led to widespread debate and proposals for the adoption of alternatives [1–3]. One moderate proposal is to switch from the reporting of *p*-values to the reporting of Confidence Intervals (CIs) [4]. This is a shift of emphasis from a dichotomous division between zero and non-zero effect size to estimating the effect size and placing confidence bounds on this estimate. CIs are based on the same underlying theory as *p*-values, providing control of the type I error probability (the probability of the false rejection of a true hypothesis) [5]. For example, Cochrane [6, section 12.4.1] uses CIs to judge whether an intervention has not just a non-zero effect but confidently a clinically useful effect. The widely used Publication Manual of the American Psychological Association [7, section 2.07] recommends giving estimated effect sizes and strongly recommends that these be accompanied by CIs, with effect sizes to be given in the original units and possibly also in a standardized form such as Cohen’s *d*.

One area this shift has not yet occurred is in differential expression analysis of microarray and RNA-Seq data. Here the effect size of interest is generally the Log2 Fold Change (LFC) in the relative RNA abundance of each gene between two groups of biological samples. A possible reason is that due to the large number of genes tested, multiple testing correction is necessary in differential expression analysis in order to maintain a False Discovery Rate (FDR) [8]. The dependence of FDR control on the number of discoveries made makes it difficult to reconcile with the use of CIs. An alternative would be to control the Family-Wise Error Rate (FWER) using a Bonferroni correction, which has a straightforward corresponding Bonferroni correction for CIs. However, unless conclusions depend on every single CI being correct, controlling the FWER is unnecessarily strict. A final possibility is a procedure that declares a certain number of discoveries made based on some criterion, and then reports False Coverage-statement Rate corrected CIs for the selected genes [9]. This has been implemented in the context of differential gene expression [10], with the criterion being that the genes have non-zero differential expression with a given FDR. An appealing feature of this approach is that the confidence intervals of genes that are just judged to be statistically significant also just touch zero LFC. However, this re-introduces a dichotomous hypothesis testing step into a CI method, where the point of using CIs is to move away from such dichotomous decisions.

Considering the current *p*-value based practice, depending on the nature of the experiment, quality of the data produced, and the chosen FDR, the number of significantly differentially expressed genes discovered may range from just a few to thousands. However, a researcher may only have resources to follow up a limited number of genes, so that thousands of discoveries present a problem. Moreover, many genes that pass this statistical test for significance may not be sufficiently changed to be of biological significance. It is important therefore to have a way to rank genes by interest level.

A common default presentation of differential expression analysis results is to list genes in order of an “adjusted *p*-value”. The researcher may choose a cutoff value for this adjusted *p*-value, producing a set of genes with that FDR. This then gives the researcher a means to select as many genes as they are able to further investigate: read down the list until the desired number of genes is obtained (with a small technicality that if multiple genes have the same adjusted *p*-value, all or none of them should be chosen). However statistical significance is not the same as a biologically meaningful effect size. It may be that in order to obtain a manageable set of genes by this method, the researcher chooses a far smaller FDR than they actually require. Genes may as easily be chosen by this method for low biological and technical variation as for a large LFC.

McCarthy and Smyth [11] propose a principled solution to this problem with their TREAT method. The researcher nominates a minimum LFC effect size of interest. The TREAT method finds genes with magnitude of effect size larger than this. Again, the researcher is presented with a list in order of adjusted *p*-value, and may make the final choice of FDR. However, how to choose the minimum effect size with TREAT is not necessarily obvious. On the other hand, the researcher may well be able to nominate an acceptable FDR (5% is a common choice).

Therefore, we describe here a new approach to the presentation of TREAT results in which the FDR is fixed, and genes are presented in order of a quantity we call the “confident effect size” or “confect”. If a set of genes is chosen having magnitude of confect greater than or equal to some amount, we guarantee with the given FDR that those genes will have a true LFC magnitude greater than that chosen amount. The researcher is then easily able to choose a desired effect size of interest to follow up, and is never presented with unreasonably small adjusted *p*-values. This is “test inversion”, converting hypothesis testing into a confidence bound calculation, however with a novel feature being the incorporation of FDR control. The confect ranking solves two problems at once, giving confidence bounds with an appropriate level of multiple-testing correction, and simultaneously providing a ranking of genes by confident effect size.

We show using synthetic data that the confect ranking method scales across experiment sizes. The method is then applied to a cancer data-set, which has a high degree of heteroscedasticity between genes. The confect ranking method, as compared to the *p*-value ranking method, leads to a markedly different emphasis on affected biological processes.

## Method

The confidence bound calculation requires as input a *p*-value function *p_i_* (*e*) for each gene *i*, 1 ≤ *i* ≤ *n*_gene_, for a test that the absolute effect size is at most *e*. *p_i_* (*e*) will be a monotonically increasing function of *e*. The TREAT method [11] provides a suitable *p*-value function, with the effect size being LFC. The `limma` R package [12] provides an implementation of this in the `treat` function.

In the next section, confidence bounds derived from TREAT are considered for a fixed significance level cutoff *α*. In the following section, this is extended to FDR control, in which confidence bounds are found with a dynamic significance level cutoff.

### Confidence bounds from TREAT

There is a close relationship between CIs and *p*-values. For example, considering the two-sided *t*-test that the LFC of a gene is not *e*, those values of *e* where *p^t^*^−test^(*e*) > *α* form a 1 − *α* confidence interval. Similarly, a confidence bound can be obtained from the one-sided *t*-test. This is called test inversion.

Note in particular that a significant result on a *t*-test for *e* = 0 not only establishes that the LFC is non-zero, but also establishes that the sign of the LFC is known, since the corresponding confidence interval will lie either entirely above or below 0.

In the case of TREAT, the null hypothesis is that the LFC lies inside the range [−*e*, *e*]. Thus TREAT *p*-values are always larger than those from the *t*-test that the LFC is 0, and a significant TREAT result determines the sign of the LFC. Taking the small liberty of considering that these two properties hold simultaneously, we view the largest *e* for which *p_i_* (*e*) ≤ *α* as providing a confidence bound, establishing either that the LFC is greater than *e*, or establishing that it is less than −*e*.

### Calculation of confects

Using TREAT, and making the assumption that each gene is independent of the others, a set of genes with effect size exceeding *e* at a given FDR *q* may be obtained using the procedure of Benjamini and Hochberg [8]. This set *S*(*e*) is the *largest* set satisfying

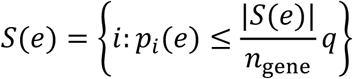

Sets for different effect sizes nest. If *e* > *e*′ then *S*(*e*) ⊆ *S*(*e*′). Genes may drop out of *S*(*e*) as *e* increases for two reasons. Firstly, *p_i_*(*e*) may rise above the threshold for inclusion in the set. Secondly, the threshold for inculsion in the set is a function of the size of the set |*S*(*e*)|, so as the set becomes smaller the threshold also becomes stricter. Thus as one gene drops out, several more may also need to immediately be dropped.

Let |*c_i_*| be the largest *e* such that *i* ∊ *S*(*e*), and let the sign of *c_i_* be the actual sign of the estimated effect. We call this quantity the “confect”, for *con*fident ef*fect* size. In our implementation, when computing *c_i_* we scan through a discrete set of effect sizes, by default considering *e* = 0, 0.01, 0.02, 0.03, … until *S*(*e*) is empty.

By presenting genes in order from largest to smallest |*c_i_*|, the researcher may easily choose an effect size resulting in a set of genes *S*(*e*) of a size suitable for their purpose. It may happen that some genes have the same |*c_i_*|, and in order to obtain a total order we sort these by *p_i_*(*e*) at the first *e* for which they are not in *S*(*e*). Some genes are not a member of any set, and are not given a confect. These are listed last, in order of *p_i_*(0) (the *p*-value given by `limma` without using TREAT). An illustration of this method is shown in Figure 1.

**Figure 1.**
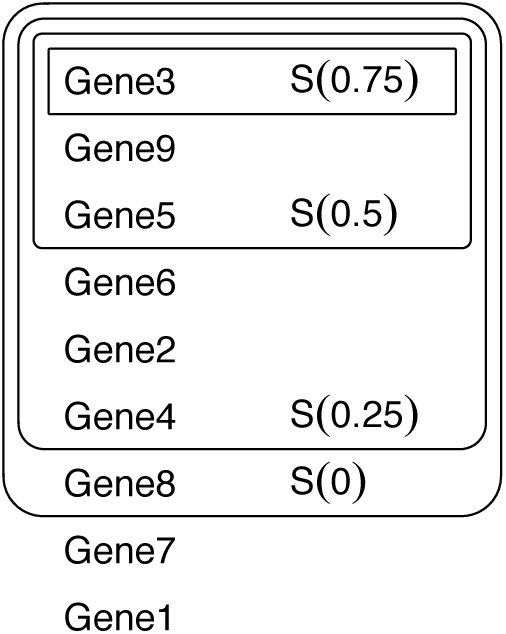
Illustration of ranking method. Sets S(e) are sets of genes with effect size significantly exceeding threshold e at some desired FDR. These sets nest, providing a ranking of genes.

The overall effect of this procedure is to provide a lower confidence bound on the magnitude of LFC for each gene, but with a higher level of confidence required for the larger effect sizes at the top of the list than for the smaller effect sizes lower down the list. Further the set {*i*:|*c_i_* | ≤ *e*} is precisely *S*(*e*), and is always at the top of the ordering.

R code implementing this procedure is provided as a supplemental file. Ongoing development of this method as an R package is available at https://github.com/pfh/topconfects

### Evaluation with synthetic data

Simulated data for *n*_genne_ genes is generated for two equally sized groups with *n*_rep_ samples within each group. We follow the distributional assumption of `limma` [12] that the gene-wise within-group variances 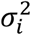 follow a scaled inverse chi-square distribution with degrees of freedom *d*_within_ and scale parameter *S*_witnin_.

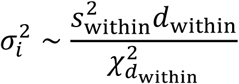

`limma`’s calculation of *p*-values, both normally and with the TREAT method, do not depend on any assumption about the distribution of LFC. `limma`’s calculation of the posterior log-odds *B* statistic does make such assumptions, specifically that there are a set of genes that are not differentially expressed, and the ratio of LFC to *σ_i_* for the differentially expressed genes follows a specific distribution. This *B* statistic is not used here. The intent of this paper is to move from the dichotomous mode of thinking associated with *p*-values to the estimation mode of thinking associated with effect sizes [4], so our simulation does not assume any gene has precisely zero LFC. However in a typical experiment some genes are differentially expressed to a much greater extent than the majority. Therefore we use a distribution with tails following a power law, specifically a scaled t-distribution with *d*_between_ degrees of freedom and scaling factor *S*_between_.

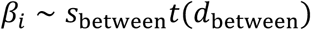

In particular the values used for the simulation were *n*_gene_ = 15000, *n*_rep_ = 2 to 10, *d*_within_ = ^2^, *S*_within_ = 0.75, *d*_between_ = 3, and *S*_between_ = 0.5. Note in particular that *d*_within_ has been chosen to be extremely small, which will generate a highly heteroscedastic data-set, in order to emphasize differences between different ways of ranking genes. Results are averaged over 100 runs of the simulation.

Eight different methods of ranking genes are compared:

a. Confect ranking at FDR 0.05. This is the proposed method, with a reasonable choice of FDR.
b. Confect ranking at FDR 0.5. This is an unreasonably high FDR, however its accuracy as a ranking method is of interest.
c. Ranking by the inner end of a 95% CI. Where the CIs span zero, genes are further ranked by `limma` *p*-value. While this does not control the FDR, its accuracy as a ranking method is of interest.
d. Ranking by the inner end of a Bonferroni corrected CI maintaining a FWER of 5%. Where the CIs span zero, genes are further ranked by `limma` p-value. This is a very strict correction for multiple testing.
e. Ranking by TREAT *p*-value with LFC threshold 1.0. While not a general ranking by LFC effect size, this should serve to distinguish genes having LFC magnitude exceeding 1.0 from those that do not.
f. Ranking by TREAT *p*-value with LFC threshold 5.0.
g. Ranking by `limma` *p*-value (i.e. TREAT LFC threshold 0.0). This is included because differential expression software often outputs genes ranked by *p*-value as the default.
h. Ranking by the magnitude of the LFC estimated by `limma`. If the noise level 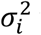 was uniform over all genes, this would be the ideal ranking method. The ranking methods that perform better than this one will do so based on their ability to adapt to heteroscedasticity.

### Evaluation with cancer data

RNA-Seq read counts for genes for 97 tumor-normal pairs from The Cancer Genome Atlas breast cancer dataset were obtained from Firebrowse [13]. There was an average of 85 million reads counted per sample. The `edgeR` R package was used to estimate TMM-adjusted library sizes [14], and the `edgeR` function `cpm` was then used to convert the count data to log_2_ Reads Per Million (RPM), using the default prior count of 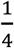. Genes with an average log_2_ RPM less than −4 were excluded from further analysis. The `limma` R package was then used to fit linear models for each gene suitable for performing a paired-samples test for differential expression between tumor and normal samples [12]. Empirical Bayes variance moderation was applied, and the `trend` option was used to allow prior variance to follow a trend line based on average expression level. The method described above was then used to calculate confect values and rank genes, using an FDR of 0.05.

### Gene-set enrichment

In order to better understand the biological processes emphasized by different methods of ranking genes, R package `fgsea` was used to find enriched gene-sets associated with Gene Ontology (GO) Biological Process terms [15]. `fgsea` implements the Gene Set Enrichment Analysis (GSEA) method, in particular the variant of the method for a pre-ranked list of genes [16]. The exponent parameter p is set to 0, so that results are based purely on the ranking, and not any associated score. The effect size used to rank gene-sets was the Normalized Enrichment Score (NES) produced by this method. A *p*-value testing whether the NES is non-zero is available from `fgsea`, but unfortunately no confidence interval. Gene-sets containing between 15 and 2,000 genes were considered.10,000 permutations were used when calculating *p*-values.

## Results

### Confect ranking out-performs alternative ranking methods in simulated data

To test the performance of the confect ranking method against alternative ranking methods, we generated simulated data-sets with between 2 and 10 replicates per group, with parameters as described in the methods section. The parameters were chosen to emphasize differences between ranking methods, and in particular the within-group variance has been made to vary greatly between genes. As the data is simulated, the true LFC for each gene and the correct ranking of genes by magnitude of LFC is known, and results from different ranking methods may be compared to this true ranking. The percentage correct genes in the top 20, 100, and 500 genes were calculated (Figure 2).

**Figure 2.**
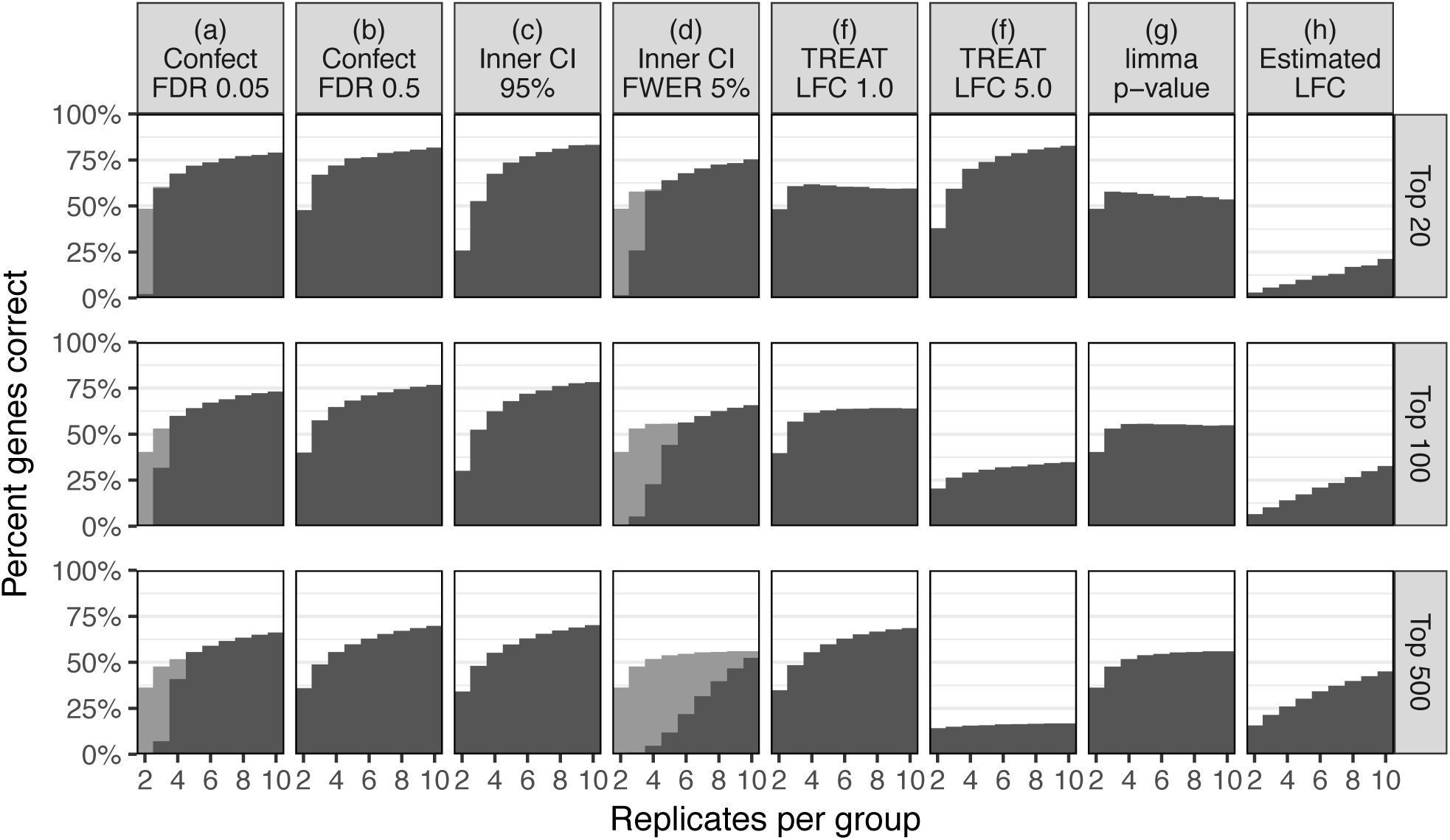
Results of simulation, showing proportion of top genes correct by various ranking methods in the top 20, 50, and 500 genes. Where genes were correct only because the ranking method fell back to ranking by `limma` p-value, this is shown in grey.

Confect ranking at FDR 0.05 (a) performs well, although for experiments with small numbers of replicates this is due to falling back to ranking by the p-value for the gene having non-zero LFC (shown as a grey bar in Figure 2). Confect ranking at FDR 0.5 (b) also performs well and without any fallback, however the confect values produced here would not be trustworthy for anything other than a method of ranking, as up to 50% of the confidence bounds within a set of top genes may be incorrect.

Interestingly, the naive method of ranking based on the inner end of a CI (c) also performed well, although performing worse than confect ranking with a very small number of replicates. The inner end of a FWER corrected CI (d) performed worse than confect ranking for larger numbers of replicates when looking for the top 100 or 500 genes.

TREAT *p*-value based ranking (e, f) may be tuned to perform well when finding a certain number of top genes, but is not a good general ranking scheme. This is as expected. The point of the confect value calculation is to modify the presentation of TREAT results to correct this shortcoming.

Although it is probably the most common approach to the analysis of differential gene expression, *p*-value based ranking (g) did not perform well in this simulation, nor should it be expected to as p-values are not an indication of LFC effect size. The estimated LFC (h) performed worst in this simulation. This is because some genes with very high within-group variability will have randomly had a large estimated LFC, displacing the genes with truly large LFC.

### In a cancer data-set, sorting by confident effect size rather than *p*-value highlights different biological pathways

To understand how ranking by confect rather than p-value impacts the interpretation of real experimental data, we turned to tumor-normal comparisons of breast cancer patients within the TCGA. With this breast cancer data-set, `limma` assigns a low prior degrees of freedom of 3.4, indicating a high degree of heteroscedasticity: different genes have very different levels of variability. The variance moderation applied here by `limma` is minor in relation to the 96 residual degrees of freedom.

Of the 17,426 genes tested, 13,416 are found to be differentially expressed at FDR 0.05 (this also means 13,416 genes are given a confect value at FDR 0.05). Such a large list is of little use to a biologist prioritizing genes for further investigation. Therefore we compared the top 20 genes ranked by confect at FDR 0.05 (Figure 3) and the top 20 genes ranked by `limma` *p*-value (Figure 4). The full rankings are included in supplemental files. The facetted plots to the right of the main listing in these figures show the raw data for each gene. The two methods of ranking have highlighted very different patterns of gene expression. Ranking by confect, the top genes have large LFC. The variability in LFC between patients is high in these genes, however the confect values are also large, giving confidence that the population average LFC is truly large. Note that sets of genes at the top of the confect ranking can be obtained using the TREAT method directly. For example, the top 10 genes would be obtained using TREAT with an LFC threshold of 5.02 (the absolute confect value for the 10^th^ gene in the ranking). However, arriving at this threshold without using confect values would require trial and error.

**Figure 3.**
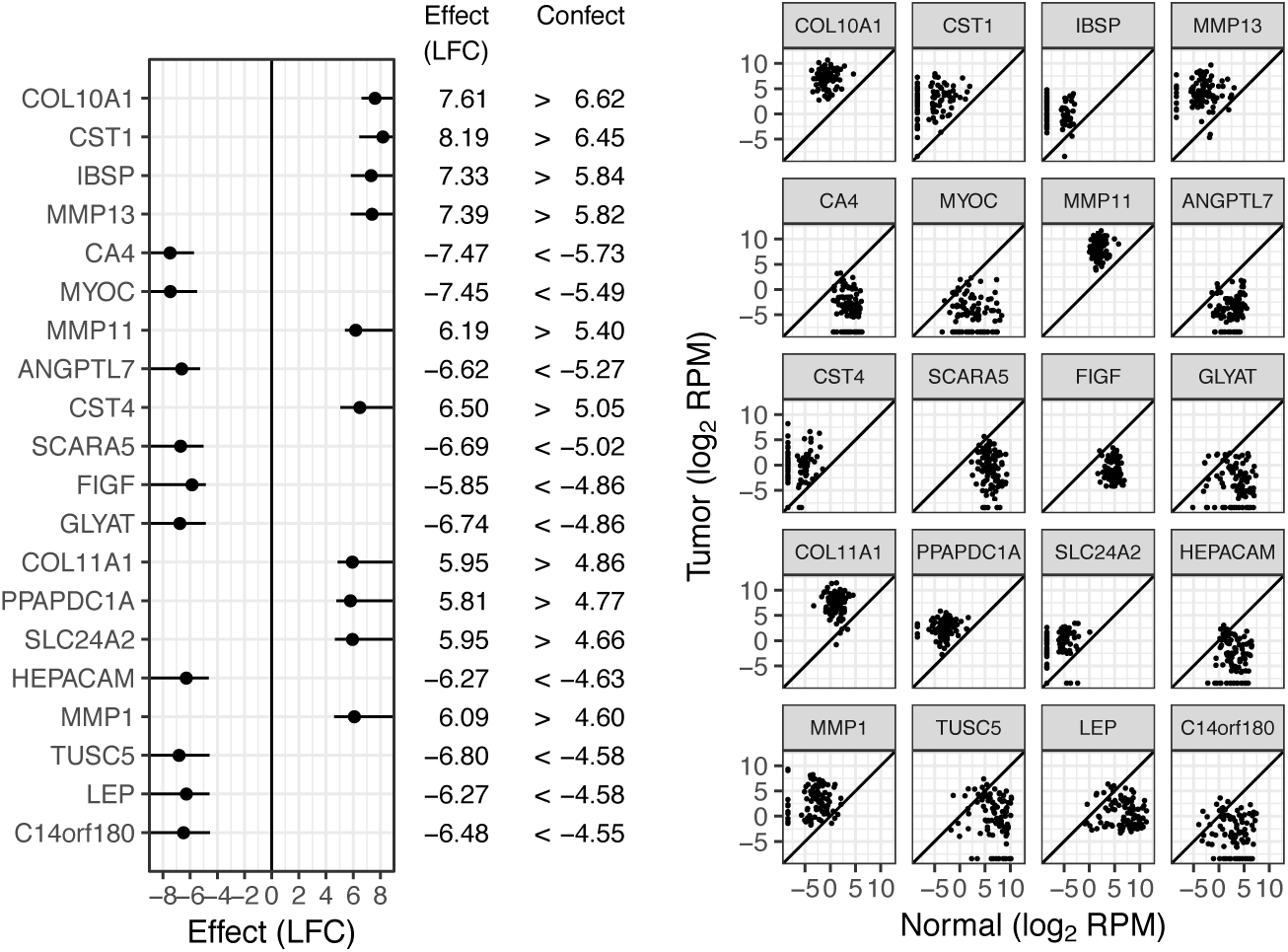
Top 20 genes by confect ranking of the breast cancer data-set at FDR 0.05. For each gene, the dot shows the estimated LFC and the line shows the “confect” confidence bound. To the right, normal and tumor expression levels for all patients are shown for each listed gene.

**Figure 4.**
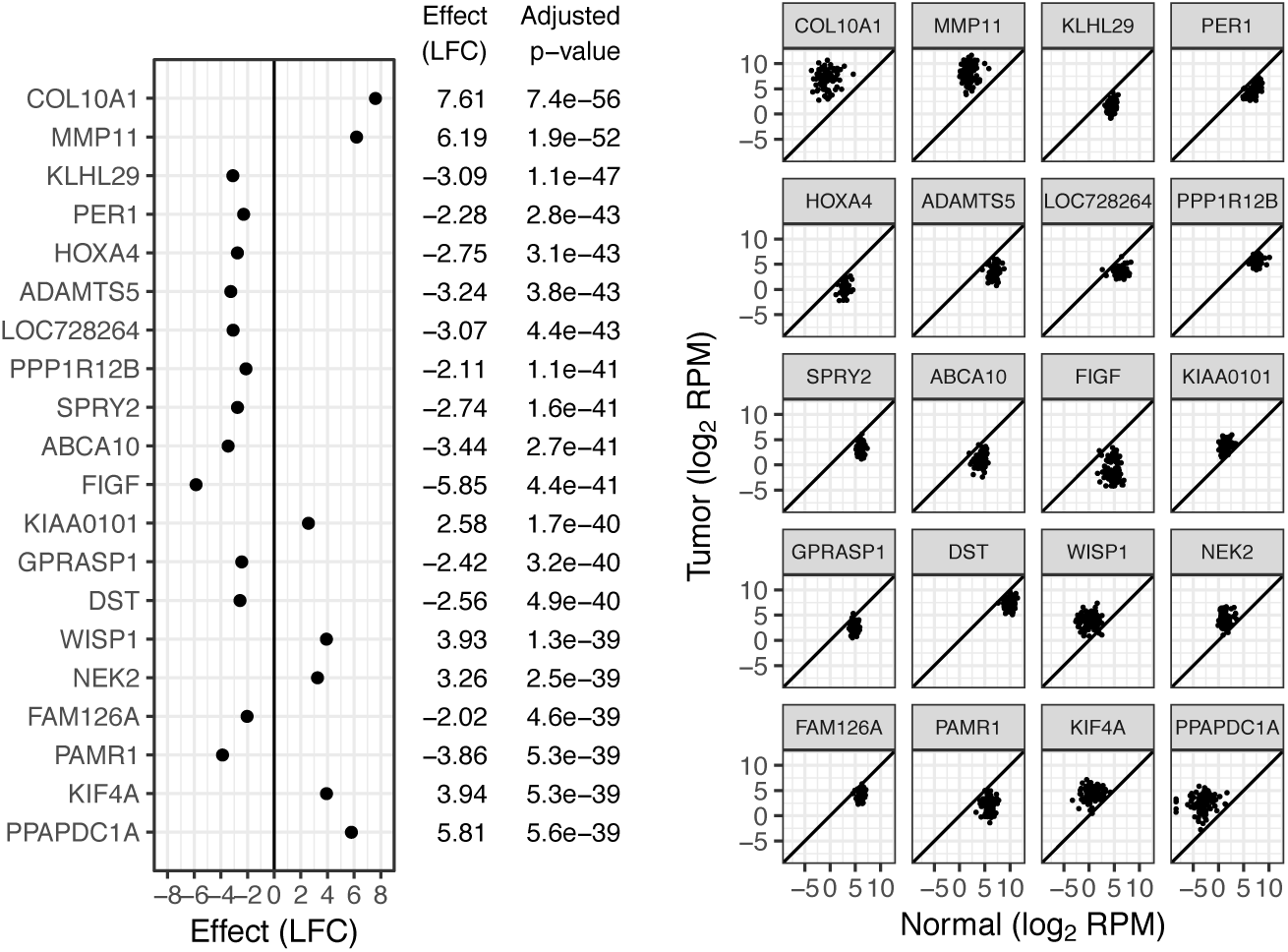
Top 20 genes by `limma` *p*-value based ranking of the breast cancer data-set. p-values shown are FDR adjusted.

Examining the ranking by *p*-value, the top genes may have smaller average LFCs if the LFC also has smaller variability between patients. Examples of such genes are NEK2 and KIF4A, both involved in chromosome segregation for cell division.

Gene-set enrichment was searched for using the R package `fgsea`. There were 5,058 GO Biological Process gene-sets available with between 15 and 2,000 genes. At FDR 0.05, 482 of these gene-sets are significantly enriched when ranking genes by p-value, and 1,336 are significantly enriched when ranking by confect. This is too many gene-sets to reasonably examine, so the Normalized Enrichment Score effect size was used to find the top enriched gene-sets. Table 1 shows the top 10 enriched gene-sets for both ranking methods. For the p-value ranking, the emphasis is on processes associated with cell division as can be expected for oncological cell transformation. For the confect ranking however, a variety of biological processes are found at the top of the list, including cell adhesion and blood vessel development suggestive of the tumor micro-environment. Also notable is the presence of genes involved in the extra-cellular matrix in the top 20 genes, including two collagen (COL10A1, COL11A1) and three matrix metalloproteinase genes (MMP13, MMP11, MMP1). Only two of these are seen in the top 20-genes by *p*-value.

**Table 1.**
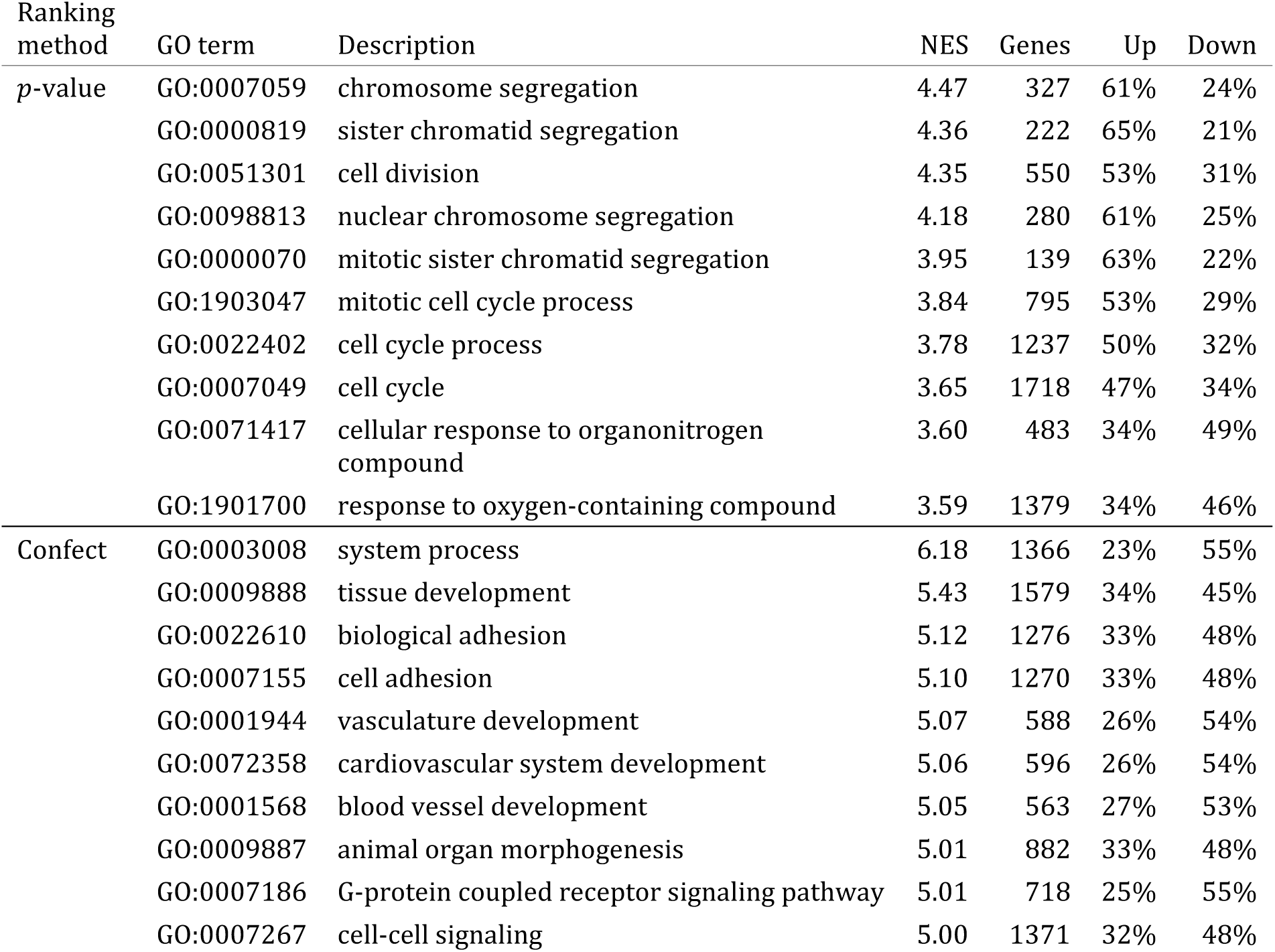
Top enriched GO Biological Process gene-sets by NES, based on p-value and confect rankings. Columns “Up” and “Down” are the percent significantly up- and down-regulated genes in the cancer samples at FDR 0.05.

Few biological experiments contain the very large number of samples present in consortia data such as the TCGA. A smaller data-set may be simulated by taking a random subset of patients. Results using a random subset of 10 patients are shown in Figure 5. For the top ranked genes, the confect values are a much smaller fraction of the effect sizes than with the full data-set. Not all of the genes with large effect sizes found in the full data-set are near the top of the list in this subset, and some genes with smaller effect sizes have been “lucky” and are highly ranked, such as SFRP1 (jumping from 53^rd^ in the full data-set to 6^th^ in the subset). “Luck” of this kind is inevitable in a small data-set with this level of heteroscedasticity, and the small confect values warn that this is occurring. Similarly, by conventional *p*-value based differential expression analysis, genes in an underpowered experiment would need a combination of a large effect size and a certain amount of “luck” to be declared significantly DE. Also note that if TREAT were being used directly, the LFC threshold would need to be adjusted between the full dataset and the subset in order to obtain a set of genes of reasonable size. The LFC threshold in TREAT may be viewed as a threshold on the confidence bound and not the effect size itself, and hence needs to be adjusted to suit the size of the experiment. The confect ranking method removes this need for parameter adjustment.

**Figure 5.**
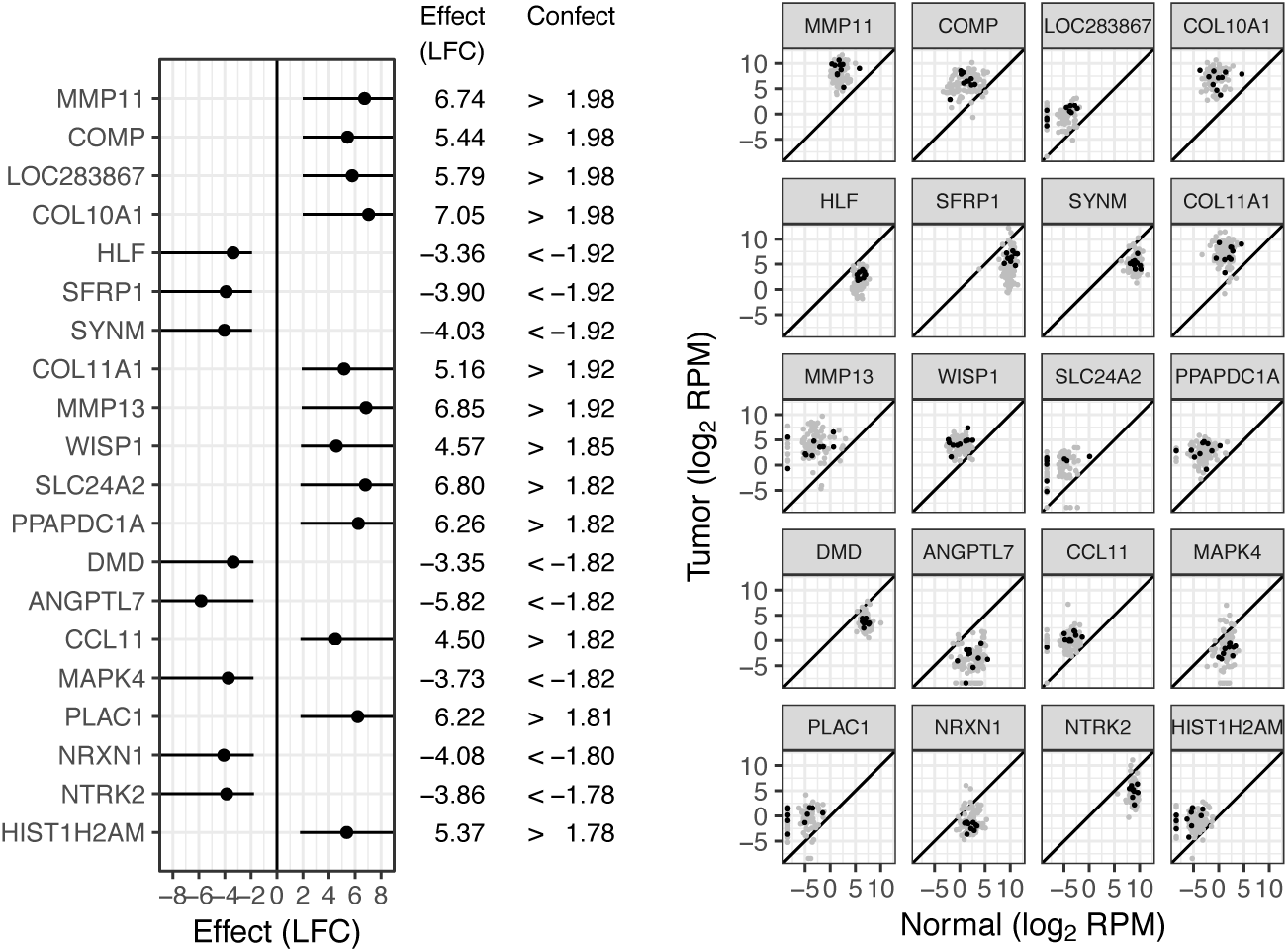
Top 20 genes by confect ranking of the breast cancer dataset at FDR 0. 05, using only 10 patients’ data.

## Discussion

The effect size used here was the LFC, with the intention of finding changes in expression with a large biological effect. The confect ranking method identifies genes with confidently large LFC. This places all genes on the same scale, and this scale has meaningful units of log2 fold change.

Can a case for using *p*-values as an effect size be made? What follows is an attempt. In fields such as psychology where the thing being measured may not have a scale with meaningful units, or where there may be a number of different scales on which something may be measured, standardized effect sizes are used. Cohen’s *d* is one such standardized effect size. Cohen’s *d* is the ratio of an effect size to some appropriate standard deviation (several choices are possible [4, 17]). Applied to differential gene expression, a problem is that each gene has its own standard deviation and is therefore effectively placed on a different scale, but a situation where comparing Cohen’s *d* between genes might be appropriate would be to identify reliable prognostic biomarkers, where the interest is in genes for which the signal exceeds the background noise level. Leaving aside the use of variance moderation in `limma`, and when the standard deviation used is the residual standard deviation of the linear model used, Cohen’s *d* is proportional to the *t* statistic, and *p* is a monotonic function of |*t*|, so *p*-values can serve as a kind of standardized effect size, albeit one that is not comparable between experiments. While the *p*-values shown in Figure 4 are meaninglessly small when considered as *p*-values, they may have some meaning when considered in this way.

In the breast cancer data-set, it was seen that different ranking methods lead to an emphasis on different biological processes, both in the top ranked genes and in downstream gene-set enrichment analysis. The difference may be largely explained by the difference in ranking between Cohen’s *d* and LFC effect sizes. The common practice of using the *t* statistic for gene-set enrichment tests is effectively a choice to use Cohen’s *d*, as discussed above.

`limma`’s TREAT method was used here as the basis of the confect calculation. The TREAT method has been extended to negative binomial Generalized Linear Models (GLMs) and Quasi-Likelihood models in the `edgeR` R package’s `glmTreat` function, specifically the `worst`. case mode [18]. The DESeq2 R package [19] also provides a test relative to a threshold for negative binomial GLMs, which could serve as a basis for the confect calculation.

It was noted in the methods section that a number of genes may fall out of *S*(*e*) simultaneously as *e* increases. This may lead to several genes being given an identical |*c_i_*| value, in which case they should ideally be included or excluded from further investigation as a group. While this is an odd feature of confect values, it is no stranger than the current practice of FDR-adjusting *p*-values. This may similarly produce a number of genes with identical adjusted *p*-values, despite having distinct unadjusted *p*-values. This is not an error but a feature of the adjustment, and again these should be included or excluded as a group.

## Conclusions

The confect ranking method described here makes best use of any amount and quality of data. There is only one parameter, the desired FDR, for which a sensible default can be given. The resulting confect quantities are used in a similar way to FDR adjusted p-values to select a set of genes of interest, and have some similar properties. However, confect values are in the same units as the effect size (here LFC), making them easier to interpret. Comparing confect values to estimated LFC values provides feedback on whether or not an experiment was under-powered. The common practice of performing an ad-hoc filtering step by estimated LFC is no longer necessary, and compared to TREAT, which provides a more principled method of filtering by LFC, even the need to provide a threshold is removed. Overall, this method of differential expression analysis has improved usability, with less expertise required in the choice of parameters and in interpretation.

## Competing interests

The authors declare that they have no competing interests.

## Author’s contributions

Paul Harrison developed the method and R code. Andrew Pattison helped test the method, and helped with the use of TCGA data. Identification of the need for a flexible means to find the most “interesting” genes from a data-set and how to go about this evolved during extensive discussions between Paul Harrison, David Powell, and Traude Beilharz.

## Acknowledgements

The results published here are in whole or part based upon data generated by the TCGA Research Network: http://cancergenome.nih.gov/

THB is supported by a Biomedicine Discovery Fellowship from Monash University, and acknowledges support the Australian Research Council (ARC Discovery Project DP170100569) and National Health and Medical Research Council (NHMRC Project Grant APP1128250).

